# Increased CD3 Immunoreactive Cells Persist Chronically in the Brain Parenchyma in Association with Focal Cortical Contusion following Experimental TBI

**DOI:** 10.64898/2026.02.13.704874

**Authors:** Kristen D. Esannason Munroe, Alexandra V. Ulyanova, Rebecca Donahue, John D. Arena, D. Kacy Cullen, Douglas H. Smith, William Stewart, Victoria E. Johnson, John A. Wolf

**Author notes:** **To whom correspondence should be addressed:** John A. Wolf, Ph.D., Department of Neurosurgery, Penn Center for Brain Injury and Repair, 172A Stemmler Hall, 3450 Hamilton Walk University of Pennsylvania, Philadelphia, PA 19104, USA. Denotes Co-Senior Authors (William Stewart, John A. Wolf). Deceased (Victoria E. Johnson).

## Abstract

While a history of TBI is associated with an increased risk of neurodegenerative disease, associated mechanisms remain largely unknown. Neuroinflammation is commonly implicated as playing a role in progressive neurodegeneration in general, yet little is known about the adaptive response of neuroinflammation in TBI or how it may contribute to progressive pathologies. To parse out components of the adaptive response, we assessed for intraparenchymal T-cell infiltration in two different translational large animal (swine) models of TBI, inertial injury and focal contusion. We characterized the extent and distribution of T cells post-injury and their association with blood-brain barrier disruption and axonal pathology. T-cell infiltration following focal TBI followed a spatiotemporal progression from gray matter at 72 hours to both gray and white matter at 6 months post-injury, consistent with recruitment into the parenchyma and then white matter. Inertial injury did not lead to substantial T-cell infiltration despite BBB breakdown and axonal pathology. We did not find a spatial correlation between blood-brain barrier breakdown or axonal pathology and T-cell infiltration in focal TBI. These data suggest that there is an active adaptive response to TBI, particularly in tissue proximal to contusions. A large animal model that reproducibly demonstrates chronic T-cell infiltration may allow for examination of the downstream effects of the adaptive response to TBI, and whether targeting this adaptive response may reduce chronic inflammation and improve recovery.

## INTRODUCTION

Approximately 10 million people annually in the United States sustain a traumatic brain injury (TBI), resulting in almost quarter of a million hospitalizations and risk of lifelong disease burden among survivors [5, 28]. Among these late outcomes, there is increasing recognition that prior history of TBI is associated with higher risk of a range of neurodegenerative disease, including Alzheimer’s disease and chronic traumatic encephalopathy [12, 14]. However, the biological processes driving an acute biomechanical injury to a prolonged, lifelong neurodegenerative disease culminating in dementia remain unknown.

Among the pathologies of TBI, neuroinflammation emerges as a likely candidate with potential to promote progressive neurodegeneration [18]. The innate immune response to TBI has been characterized in both animal models and human TBI, with emerging evidence of primed microglial activation leading to a more chronic, persistent activated state [11]. In contrast, the adaptive immune response to TBI remains unclear, as does the potential interaction between the adaptive and innate responses over chronic time periods [32]. T-cells in particular have been demonstrated to infiltrate the brain parenchyma in both rodent models and human TBI. The primary evidence for their presence in human TBI comes from examination of resected contused brain tissue removed in the setting of neurosurgical decompression [9]. These findings have been difficult to reproduce using post-mortem tissue, potentially due to the transient T-cell response or their sparse nature post-injury. Nonetheless, a recent report suggests T-cell infiltration into brain parenchyma in chronic TBI [4]. In addition, autoantibody studies in both acute and chronic TBI have demonstrated substantial response to brain proteins associated with T-cell activity [21]. Extensive rodent TBI work examining the adaptive immune response has demonstrated both acute and chronic T-cell infiltration post-TBI and has even correlated these findings with recovery [19]. In addition, manipulation of the T-cell response has been demonstrated to affect chronic white matter degeneration up to 8 months post-injury, and anti-CD3 antibodies have induced regulatory T-cells that are beneficial post-TBI [6, 22]. However, the general conclusion has been that both beneficial and detrimental aspects of the adaptive response occur in parallel, and the contributions of T-cells to these processes are unclear [20, 23].

To assess the clinical relevance of rodent studies in this area, we examined intraparenchymal T-cell infiltration in two different translational large animal (swine) models of TBI, one with pure rotational acceleration to induce inertial TBI (RAI), and one producing a contusion with diffuse injury absent inertial injury, controlled cortical impact (CCI) [10]. The CCI model induces focal contusion, while the RAI model produces diffuse axonal injury and multifocal blood brain barrier disruption in the absence of hemorrhage or other gross pathologies [2, 16]. Identification of persistently infiltrating T-cells in these models may help elucidate their evolving role in after TBI in context with other persisting pathologies.

We hypothesized that T cells may infiltrate the brain parenchyma due to vascular compromise either through breaches caused by breakdown in blood brain barrier following inertial TBI, or enter through sites of overt hemorrhage following contusion. In addition, we predicted that infiltrating T-cells would overlap with regions of BBB breakdown or be guided to regions of axonal pathology. We utilized immunohistochemistry to characterize the extent and distribution of T cells post-injury and their association with specific neuropathological features of TBI, including blood-brain barrier disruption and axonal pathology.

## MATERIALS AND METHODS

### Experimental Design

All animal experiments were conducted in accordance with protocols approved by The University of Pennsylvania Institutional Animal Care and Use Committee and in accordance with the NIH Guide for the Care and Use of Laboratory Animals. Archival brain tissue from 6-8 month-old castrated male Yucatan miniature swine were examined following the controlled cortical impact (CCI) model or rotational acceleration injury (RAI) models of TBI as described below. Animals were assigned to either a 72-hour survival group (RAI, n=3; CCI, n=3; sham CCI, n=2) or a 6-month survival group (CCI, n=3; sham CCI, n=1. All animals underwent a standardized neurologic examination in collaboration with facility veterinarians post-injury to assess gait, tone, and reflexes. These assessments were formalized and documented for the 6-month cohort at baseline, and 24, 48, and 72 hours post-injury.

### Controlled Cortical Impact (CCI) Model

Following induction of anesthesia with 0.4mg/kg midazolam IM and 5% inhaled isoflurane, animals were intubated, and anesthesia was maintained with 2.5% inhaled isoflurane. Animals were ventilated and hemodynamically monitored throughout the procedure. Under sterile conditions, bupivacaine (0.25%) was administered locally to the incision site. The head was fixed in position using a custom-designed stereotaxic frame, as described in detail previously [27] and an 18 mm-diameter burr hole was drilled over the left hemisphere. For sham and 72h survival animals, the burr hole was placed 9mm lateral to the midline sagittal suture and 13mm anterior to bregma. Animals that survived for 6 months had a slightly posterior impact location at 9mm lateral to the midline sagittal suture and 4.5mm posterior to bregma. In all CCI animals, a cortical impact was performed vertically onto the intact dura using a metal impactor with a beveled 12mm diameter tip. The impact was driven by the release of pressurized nitrogen to a depth of 9mm at an average velocity of 3.54ms^-1^ and an average dwell time of 209ms (CCI device model AMS-201: AmScien Instruments, Richmond, VA). Following impact, the cranial defect was repaired using a low-toxicity silicone elastomer (Kwik-Sil^TM^, World Precision Instruments, Sarasota, FL), and the incision was closed with a running monofilament suture. Sham CCI animals underwent all procedures including craniotomy with the exception of the cortical impact. Following the procedure, inhaled anesthesia was withdrawn, and analgesia was provided post-injury in the form of 0.15mg/Kg of Buprenorphine SQ (slow-release preparation).

### Rotational Acceleration Injury Model

As previously described in detail [15, 16, 30], animals were induced using 0.4mg/kg midazolam IM and maintained via 2.5% inhaled isoflurane. The HYGE pneumatic actuator was used to induce rapid head rotation, creating linear motion by triggering the release of pressurized nitrogen. Linear motion is then converted into angular motion via custom-designed linkage assemblies to induce rotation [25]. Rotational kinematics were recorded using angular velocity transducers (Applied Technology Associates) mounted to the linkage sidearm coupled to a National Instruments data acquisition system running custom LabVIEW software (10kHz sampling rate). We produced pure impulsive centroidal head rotation of up to 110 degrees in the coronal plane with a mean peak angular velocity of 236 rad/s [3]. Animals were recovered from anesthesia and returned to the housing facility. All animals received preemptive analgesia post-injury in the form of 0.1-3mg of Buprenorphine (slow-release preparation) SQ and acetaminophen 50mg/kg PR.

### Tissue Handling and Histological Procedures

At the study endpoint, all animals were deeply anesthetized and transcardially perfused using chilled heparinized saline, followed by 10% neutral buffered formalin (NBF). Brains were post-fixed for 7 days in 10% NBF, sectioned into 5mm blocks in the coronal plane throughout the entire rostro-caudal extent of the brain, and processed to paraffin using standardized techniques [15, 16].

#### Immunohistochemical Labeling

All immunohistochemical (IHC) examinations were performed using 8µm formalin-fixed and paraffin-embedded (FFPE) whole brain sections in the coronal plane at the brain level at the epicenter of the cortical impact. Additional sections remote from the cortical impact at the level of occipital cortex were also examined for CCI animals that survived for 72h. Immunoenzymatic double labelling was performed to allow for co-examination of CD3+ cells and fibrinogen as a marker of BBB permeability [16]. Briefly, following deparaffinization and rehydration, tissue sections were immersed in aqueous hydrogen peroxide (15 minutes) to quench endogenous peroxidase activity. Antigen retrieval was performed in a pressure cooker by immersion in a preheated Tris-EDTA buffer. Subsequent blocking was achieved using Bloxall for 10m (Vector Labs, Burlingame, CA) followed by 1% normal goat serum (Vector Labs, Burlingame, CA) in Optimax buffer (BioGenex, San Ramon, CA) for 30-minutes. Incubation with CD3 (clone CD3-12, 1:500) (Bio-Rad Laboratories, Hercules, CA) was performed overnight at 4^°^C. After rinsing, sections were incubated with ImmPRESS-AP Anti-Rat Detection Kit (Vector Labs, Burlingame, CA), and detection was performed with Vector Red (Vector Labs, Burlingame, CA). Following rinsing, sections underwent incubation for 30 minutes with 1% normal horse serum (Vector Labs, Burlingame, CA) in Optimax buffer, followed by Incubation with antibody specific for full-length fibrinogen (Fibrinogen Beta, D-4, 1:20K) (Santa Cruz Biotechnologies, Dallas, TX) performed overnight at 4^°^C. After rinsing, subsequent incubation with the relevant species-specific biotinylated antibody at 1:350 was performed followed by the avidin-biotin complex (Vector Labs, Burlingame, CA) and 3,3’-diaminobenzidine DAB kit per manufacturer’s instructions (Vector Labs, Burlingame, CA). Cover slipping was performed using Vectashield mounting medium (Vector Labs, Burlingame, CA).

Single Label APP Immunohistochemistry: Serial sections also underwent single immunohistochemical labeling for APP. Specifically, following dewaxing, rehydration, hydrogen peroxide immersion and antigen retrieval using heat and pressure as above an antibody specific for the amyloid precursor protein (APP) (Millipore, Billerica, M; 1:80k) was applied overnight [7, 13, 15, 24]. After rinsing, sections were incubated with the relevant species-specific biotinylated secondary antibody for 30 minutes (Vector Labs, Burlingame, CA), followed by the avidin-biotin complex (Vector Labs, Burlingame, CA). Finally, visualization was achieved using DAB (Vector Labs, Burlingame, CA), and counterstaining with hematoxylin was performed.

Positive control tissue for APP included sections of swine tissue with previously established DAI [15]. Positive control tissue for fibrinogen extravasation included a section of swine brain tissue with previously established contusional injury with BBB permeability. Positive control tissue for CD3+ cells included a section of porcine spleen tissue. Omission of primary antibodies was performed on the same material to control for non-specific binding for each individual IHC experiment.

### Analysis and Quantification of Neuropathological Findings

#### Quantification of CD3+ Cell Infiltration

Whole-brain sections at the epicenter of the CCI, stained for both CD3 and fibrinogen, were scanned at 20X magnification (Aperio). Anatomically defined regions in both gray and white matter were examined. Specifically, gray matter regions of 960,000µm^2^ were selected spanning the impacted gyrus at the most superior aspect of the cortex, the depths of cortical sulci, and the midpoint of the gyrus. Three regions of equal size were also selected in the white matter directly beneath the sulcal depth. Corresponding regions were selected in the contralateral hemisphere. The number of CD3 immunoreactive cells was then quantified per unit area (mm^2^) in all regions. Statistical analysis was performed using ANOVAs with Tukey’s multiple comparisons test comparing the presence per unit area in regions at the level of injury, ipsilateral and contralateral to the impact site, as well as in peri-contusional gray and white matter.

#### Assessment of the Relationship Between CD3+ Cells and Vascular Compromise

To determine whether BBB permeability outside the setting of contusion was associated with increased CD3+ cells in the parenchyma, we also examined brain regions remote from the contusion at 72h post CCI, demonstrating BBB permeability. Specifically, regions immunoreactive for extravasated fibrinogen were outlined in the hemispheres, both ipsilateral and contralateral to the CCI lesion, in a region 1mm posterior to the level of the contusion in the brain. The number of CD3+ cells within these regions was quantified. The concentration of CD3+ cells within regions with fibrinogen extravasation was compared to regions without extravasation within the same tissue section using the unpaired two tailed TTEST.

#### Associations of T-Cells with Axonal Degeneration

Pathology APP-stained sections at the epicenter level were reviewed for the presence of axon degeneration, as identified by swollen varicose axonal profiles and axonal bulbs as previously described[7, 11, 24]. Briefly, injured axons were identified and defined as those that were APP positive axonal profiles with an injured morphology including 1. Terminal axonal swellings or axonal bulbs, formerly known as retraction balls. 2. Axons with a beaded or fusiform morphology representing multiple points of apparent transport interruption as extensively described previously [7, 8, 11, 15, 16, 24, 26]}. The presence or absence of pathology was recorded within the level of injury, ipsilateral and contralateral to the impact site, as well as in peri-contusional gray and white matter.

## RESULTS

### Clinical and gross neuropathological findings

Immediately following injury and while under anesthesia, no average changes in heart rate or BP were observed. Upon withdrawal of anesthesia, animals regained consciousness (mean = 25 minutes) with no evident difference in recovery time between RAI or CCI. Neurological examination revealed minor deficits in two of three CCI animals in the 6-month cohort at the 24 and 48-hour timepoints, including unstable gait and an inability to circle to the right. These resolved by the 72-hour time point. There was no evidence of acute seizure as determined by clinical observation at any time point in any animals.

Examination of the formalin-fixed whole brains at 30m and 72h survival timepoints from all animals undergoing CCI revealed evidence of both intra- and extra-parenchymal hemorrhage. Specifically, expected trace amounts of epidural blood were observed immediately beneath the burr hole site in both sham and injured animals. In contrast, subdural and subarachnoid blood was observed only in CCI animals within the impact region, was < 1 cc in volume, and showed no significant mass effect. Interestingly, blood in the subdural space was also observed in all CCI animals in the dorsal aspect of the hemisphere contralateral to the impact. This was again minimal (< 1 cc) and without significant mass effect. Finally, a minimal trace of subarachnoid blood was observed in the most posterior aspect of the cerebellum. The ventral surface of the brain was normal in all animals. On examination of blocked coronal slices, cortical contusion was observed at all time points post-injury in the region directly below the impact site, with involvement of the cortex and extension into the white matter. While not observed at 30 minutes post-CCI, by 72 hours, there was evidence of brain swelling with a midline shift at the coronal level of impact, compression of the lateral ventricles, and flattening of the gyri, most pronounced in the ipsilateral hemisphere. By 6 months post-CCI, consistent with observations acutely, there was evidence of chronic subdural blood in the cortical surface, both ipsilateral and contralateral to the impact, as well as an evolving focal cortical contusion in the aforementioned region directly under the impact site. In addition, all brains demonstrated a degree of diffuse atrophic change, maximal at the impact site, with evidence of widening of the sulci. Enlargement of the lateral ventricles was also observed maximally in the ipsilateral side.

The brains of all RAI animals appeared grossly normal without evidence of any subdural, subarachnoid, intra-parenchymal hemorrhage or other focal lesions. There was no flattening of the sulci or midline shift, and ventricles were of normal size and symmetrical in these or Sham CCI animals.

### Following CCI, there is a marked increase in peri-contusional CD3+ cells in the brain parenchyma that persists for 6 months post-injury

#### CCI Model

While there was no increase in CD3+ cells at 30 minutes (p>0.99), a marked increase in CD3 immunoreactive cells was observed in the brain parenchyma in the peri-contusional gray matter by 72 hours following CCI injury versus sham (p=0.0001) **(Fig. 1B) (Fig.3A)**. This was notably also observed to be persistent at 6 months post-CCI (p=0.0005) **(Fig.3A)**. Cells were observed diffusely in the peri-contusional region of the impacted gyri, both diffusely and with superimposed clusters. Cells could also be observed in the perivascular space and in some cases traversing the vascular boundaries into the parenchyma.

**Figure 1:**
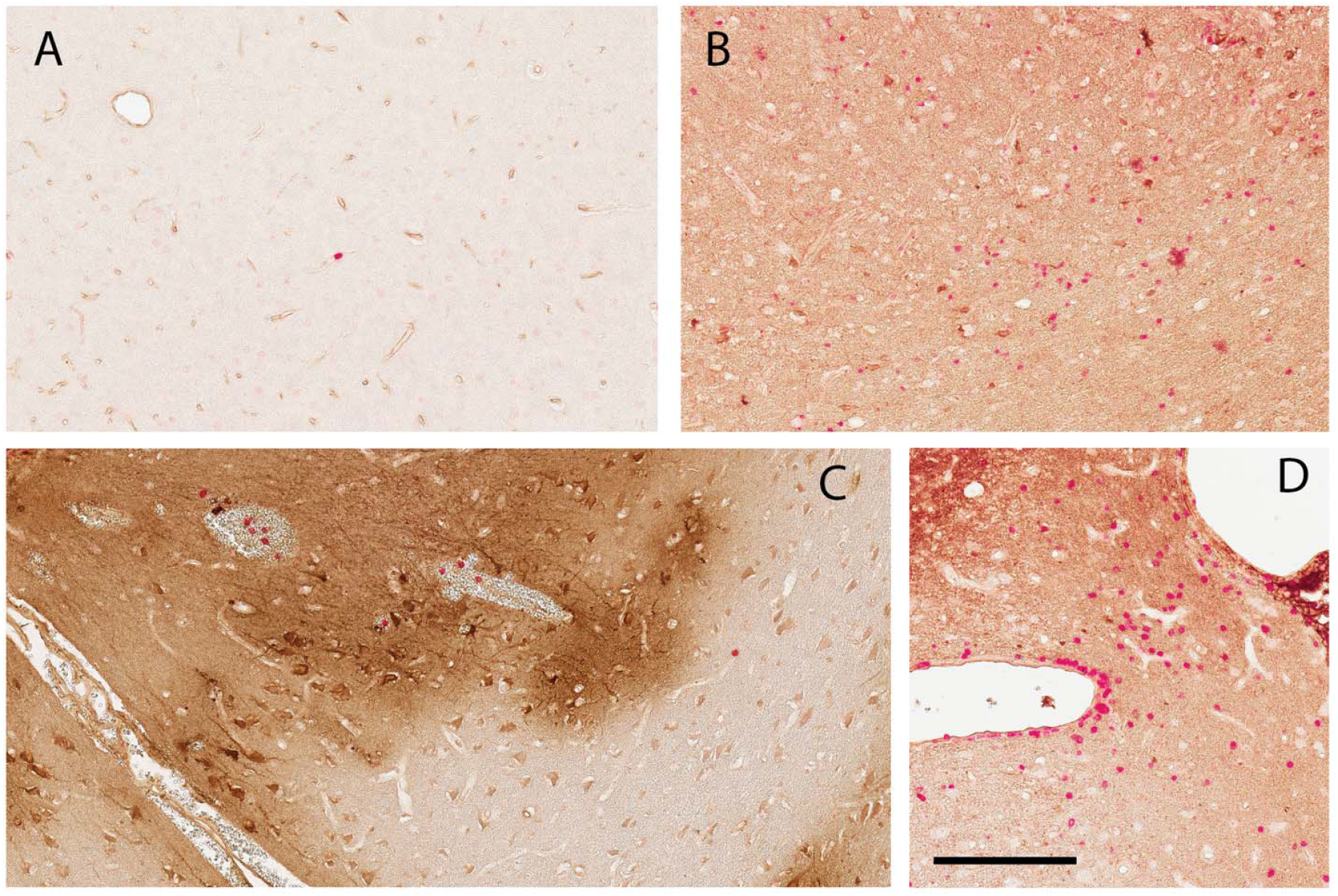
Blood Brain Barrier Breakdown and Acute T-Cell responses to Controlled Cortical Impact in the Gray Matter: A-D.) Double immunoenzymatic labeling for fibrinogen (DAB; Brown) and CD3+ cells (Vector Red; Pink). **A)** Sham showing a single CD3+within the vascular compartment. **B)** Multiple CD3+ cells in peri-contusional tissue with associated fibrinogen immunoreactivity at 72 hours. **C)** Fibrinogen extravasation indicative of early BBB permeability at 30 minutes post-CCI without overt CD3+ cell infiltration. **D)** CD3+ cells and fibrinogen extravasation, including abundant CD3+ cells around a vessel in the peri-contusional region at 72 hours. Scale bar = 200µm.

There was notably no overt increase in CD3+ cells in the white matter underlying the contusion at either 30 minutes (p>0.99) or 72 hours (p=0.9995) post-CCI, despite overt and extensive APP immunoreactive axonal degeneration in the same regions **(Fig. 3B)**. However, a marked increase in CD3+ cells was observed in this region at 6 months following CCI (p=0.012) **(Fig. 2C, 3B)**. Cells were again observed diffusely and in clusters throughout the digitate white matter and in the white matter underlying the impacted gyri **(Fig. 2B, C)**.

**Figure 2:**
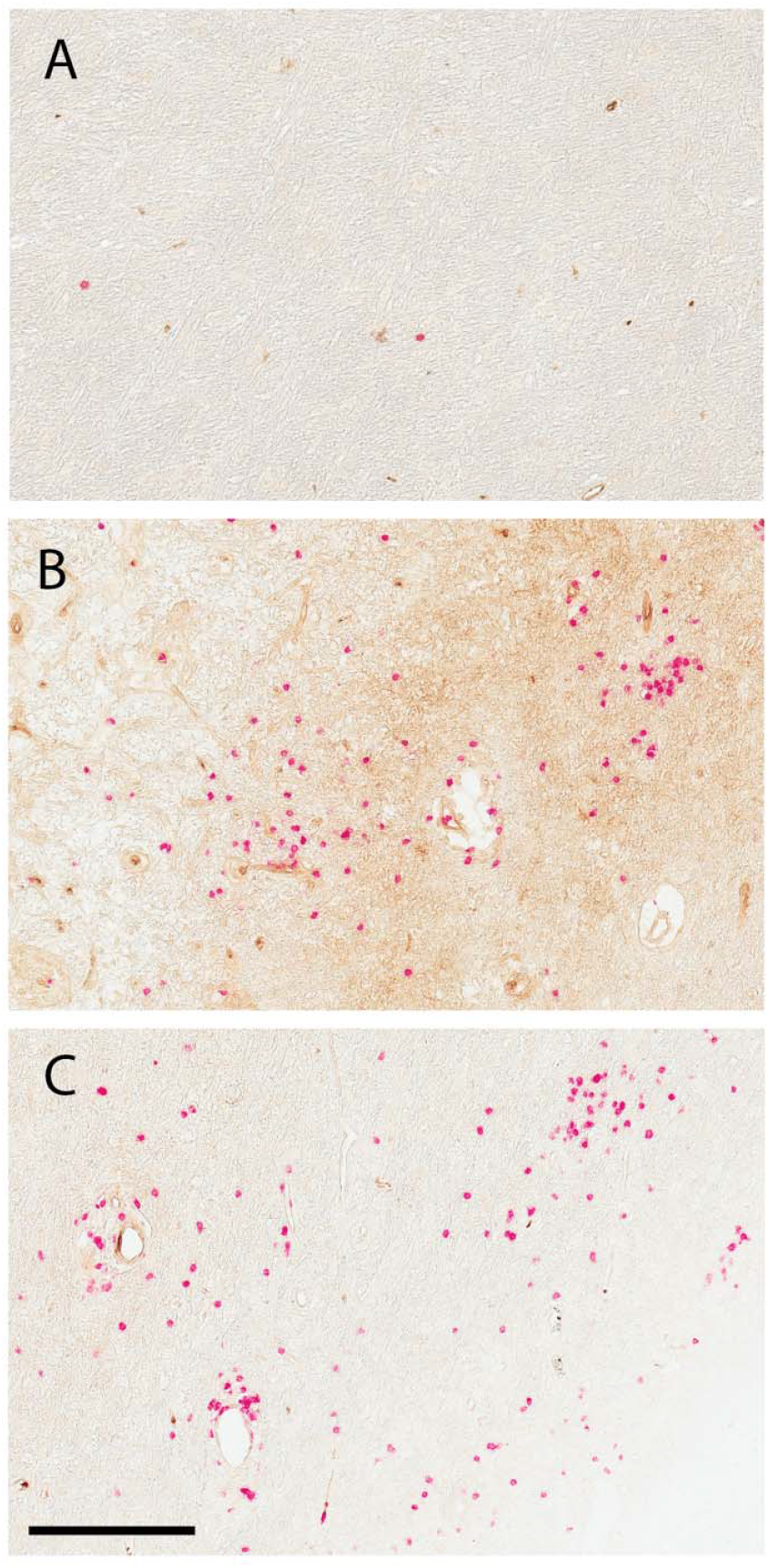
Blood Brain Barrier Breakdown and Chronic T-Cell responses to Controlled Cortical Impact in the White Matter. **A)** Minimal CD3+ cells and fibrinogen extravasation in sham at 6 months versus **B)** extensive CD3+ cells below the CCI site at 6 months in association with ongoing blood-brain barrier permeability and **C)** CD3+ cells in the underlying white matter. Scale bar = 200µm.

**Figure 3:**
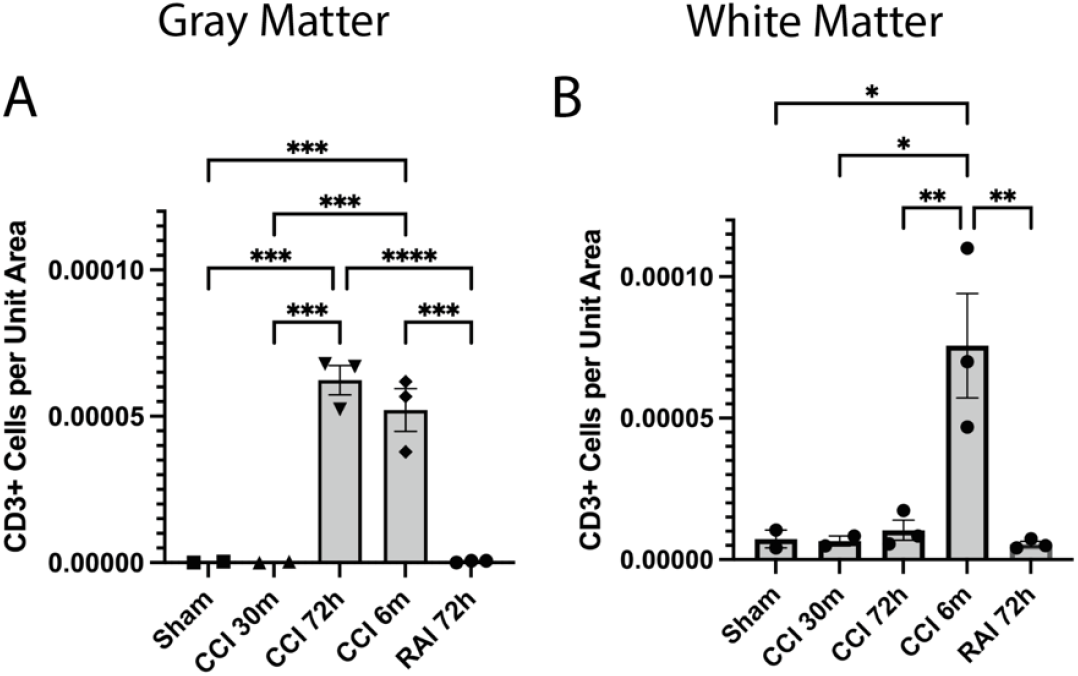
T-cells infiltrate both the gray and white matter at chronic time points following contusion. Graph showing the extent of CD3+ cells in the acute and chronic CCI model within the gray **(A)** and white **(B)** matter. There is a profound increase in CD3+ cells in the peri-contusional gray matter at the 72-hour and 6-month time points, while CD3+ cells infiltrate the peri-contusional white matter only at the 6-month time point. (ANOVA, *p= <0.05, **p=<0.01, ***p= <0.001, ****p=<0.0001).

There was no increase in CD3+ cells in the contralateral hemisphere, sampled within the same regions following CCI. Specifically, there was no increase in the contralateral gray matter at 30m (p=0.998), 72h (p<0.99), or 6 months (p<0.99) or in the contralateral white matter at 30m (p=0.64), 72h (p=0.97), or 6 months (p=0.97). Thus, increases in CD3+ cells were observed only in association with the cortical contusion on the ipsilateral hemisphere.

#### RAI Model

Notably, there was no increase in CD3+ cells versus shams following the rotational acceleration model of injury acutely at 72h in the ipsilateral gray or white matter (both P>0.99). Similarly, there was no increase in CD3+ cells compared with shams following the rotational acceleration model of injury at 72 hours in the contralateral gray matter (p=0.72) or white matter (p=0.77) **(Fig. 3A and B)**.

### CD3+ Cells in the Brain Parenchyma are associated with Contusion and not BBB Permeability Alone

CD3+ cells were notably not observed in association with BBB permeability alone. Specifically, at 72h post-CCI, when examining for regions remote from the contusion with and without fibrinogen extravasation, there was no difference in the extent of CD3+ cells in either the ipsilateral (TTEST p=0.32) or contralateral (TTEST p=0.42) hemisphere. The lack of significance in the correlations between CD3+ cells and fibrinogen in these regions suggests that contusion plays a unique role in the increase in CD3+ cells and is not spatially associated with BBB permeability. This finding suggests active recruitment as opposed to passive leakage from the vasculature due to BBB breakdown.

## DISCUSSION

The adaptive immune response following TBI may be an important mediator of long-term neurodegeneration and chronic deficits. Here, we utilized two clinically-relevant large animal models of TBI to examine T-cell infiltration into the brain parenchyma at acute and chronic time-points post injury. Our findings demonstrate that T-cell infiltration following focal TBI follows a specific temporal and spatial pattern, suggesting a progression from gray matter early to both gray and white matter at 6 months post-injury. As expected, T-cells did not appear in the contused gray matter immediately following injury but rather appeared by the 72-hour time point, consistent with recruitment into the parenchyma. Contrary to our expectations, we did not find a spatial correlation with blood-brain barrier breakdown or axonal pathology, suggesting that this process is likely mediated by other factors. In addition, inertial injury alone (RAI) did not result in increased T-cell infiltration despite extensive BBB breakdown and axonal pathology [16].

These data suggest that there is an active and persistent adaptive response of T-cells after TBI in association with contused tissue. The delayed appearance of T-cell in the CCI model suggests the contused tissue milieu mediated by active recruitment. Potentially, peripheral priming by CNS antigens drove a secondary wave of T-cell recruitment and infiltration into the parenchyma. Further support for a more targeted response comes from the finding that CD3+ cell counts were not increased in regions remote from the cortical contusion in either the acute or chronic CCI groups, despite areas exhibiting fibrinogen extravasation from vessels, suggesting blood-brain barrier breakdown. Prior work from others in our group has demonstrated changes in the peripheral immune system following two different injury severities in another plane of rotation (sagittal), including upregulation of T-cells at 14 days post injury [29]. This suggests there may be a peripheral upregulation of T-cell activity sometime between 72-hour and 6-month post-injury. However, the lack of an increase in circulating cytokines in this model suggests that recruitment of T-cells was modulated by focal aspects within and nearby the contusion site.

The marked increase in CD3+ T cells in the white matter at 6 months post-CCI demonstrates that this form of inflammation persists chronically, similar to post-mortem observations in human TBI [11]. Surprisingly, T-cell infiltration did not appear to be related to either BBB disruption or APP+ swollen axonal profiles in the CCI model. Accordingly, it appears that there remains an unknown mechanism that recruits CD3+ cells selectively into the brain parenchyma into and around contusions.

These findings in clinically relevant large animal models of TBI appear to corroborate observations in rodent models of TBI, where both acute and chronic infiltration of T-cells were identified. They also are consistent with findings in clinical studies that identified T-cells in parenchymal tissue resected from decompressive craniectomies at early time points post-injury. While the recruitment of these cells remains poorly understood, it has been posited that T cell infiltration into the parenchyma may occur via the lymphatic system via specific meningeal lymph vessels [17, 31].

Notably, T-cells in post-mortem tissue are notoriously difficult to detect, potentially due to their relatively sparse nature as well as challenges in antigenic retrieval for T-cell epitopes. Recent studies on human serum after TBI have shown a significant presence of autoantibodies to brain proteins [21]. While these studies do not directly demonstrate T-cell involvement, they strongly suggest a role for T-cells in the adaptive immune response. Extensive rodent work in this area has demonstrated T-cell infiltration at both acute and chronic time points (for review, see [20]). Examination of T-cells in rats following a fluid percussion injury noted T-cell infiltration in the perilesional cortex out to 90 days post-injury, and also concluded that T-cell infiltration did not co-localize with BBB breakdown [19]. Interestingly, despite the severity of the injury and the proximity of the corpus callosum, they did not identify T-cells in the white matter underlying the injured cortex at the time points examined, in contrast to our findings.

In a previous study of mouse CCI, a dual nature of the T-cell response was identified where subtypes of γδ T cells were found to promote pathology by activating microglial cells via IFN-γ and IL-17, whereas other subtypes promoted a homeostatic microglial signature [1]. Other studies have targeted the CD8+ cells post TBI, demonstrating that depletion of these cells in comparison to CD4+ cells leads to recovery of function [6], while stimulation of regulatory T cells using anti-CD3 antibodies has also been demonstrated to affect lesion size and function following contusion [22]. Therefore, further identification of the T-Cells in this gyrencephalic model is important future work.

There are limitations to this work, including the inability to fully phenotype the T-cells. We did not examine tissue from rotational injury at the 6-month time-point, although the lack of response at 72 hours suggests an increase after this time point is unlikely. We also did not directly quantify microglial activation, although serial sections contain activated microglia in the same regions. While a few CD3+ T-cells were noted in the parenchyma of sham animals, this is likely due to resident T lymphocytes or the effects of craniotomy [17]. We also have not examined the time points between 72 hours and 6 months, so it is difficult to know whether the time course of the infiltration is continuous or multi-modal. Understanding the phenotypic characterization of these T cells will also help further elucidate whether they are resident T lymphocytes or actively being recruited to the gray and white matter.

An important unresolved question is whether these T-cells are now “sequestered” in the white matter, creating a persistent state, or whether they are continually recruited from the periphery. If these are CD8+ T-cells, they may contribute to maintaining microglia in a chronically active state, potentially leading to chronic axonal loss, as demonstrated in human pathology and mouse models [6, 11]. Future work examining the status of the peripheral T-cell pool at later time points, where we found continued infiltration in the white matter in the CCI model, might also be informative. In the gyrencephalic brain, the high volume of white matter provides a large reservoir for lingering inflammation. Our model may provide a “missing link” between the informative rodent work and the human pathology, modeling chronic white matter inflammation following TBI. In addition, it raises the important question of whether targeting this chronic “adaptive neuroinflammation”, particularly following contusion, might be a useful therapeutic strategy for chronic neurodegeneration following TBI, as has been demonstrated in rodent models.

## Funding

Research reported in this publication was supported by:

(DoD) ERP W81XWH-20-1-0901 (JAW)

(DoD) ERP W81XWH-20-1-0838 (VEJ)

(DoD) ERP IDA W81XWH-22-1-0287 (AVU)

(VA) 5I01RX003498 (JAW)

(VA) I21 RX004629 (AVU)

(NIH) R01NS123034

(VEJ) (NIH) T32NS043126 (DHS, JA)

(DoD) PROCEED-TBI: HT9425-23-1-1039 (DHS)

(NIH) TReND: R01NS094003 (DHS)

## Acknowledgements

The content is solely the responsibility of the authors and does not necessarily represent the official views of the National Institutes of Health, the U.S. Department of Veterans Affairs, or the Department of Defense.

This work is dedicated to the memory of Victoria E. Johnson, MBChB, Ph.D.

## REFERENCES

1 Abou-El-Hassan H, Rezende RM, Izzy S, Gabriely G, Yahya T, Tatematsu BK, Habashy KJ, Lopes JR, Oliveira GLVd, Maghzi A-H et al (2023) Vγ1 and Vγ4 gamma-delta T cells play opposing roles in the immunopathology of traumatic brain injury in males. Nature communications 14: 4286 Doi 10.1038/s41467-023-39857-9

2 Arena JD, Smith DH, Arrastia RD, Cullen DK, Xiao R, Fan J, Harris DC, Lynch CE, Johnson VE (2024) The neuropathological basis of elevated serum neurofilament light following experimental concussion. Acta Neuropathologica Communications 12: 189 Doi 10.1186/s40478-024-01883-z

3 Browne KD, Chen XH, Meaney DF, Smith DH (2011) Mild traumatic brain injury and diffuse axonal injury in swine. J Neurotrauma 28: 1747–1755 Doi 10.1089/neu.2011.1913

4 Calderazzo SM, Stein TD, Holtzman DM, McKee AC, Huber BR (2024) Meningeal and infiltrating T-cells are associated with neuroinflammation after head injury and with tau pathology in CTE. Alzheimer’s & Dementia 20: e089160.Doi 10.1002/alz.089160

5 CDC (2024) Traumatic Brain Injury and Concussion https://www.cdc.gov/traumatic-brain-injury/data-research/index.html

6 Daglas M, Draxler DF, Ho H, McCutcheon F, Galle A, Au AE, Larsson P, Gregory J, Alderuccio F, Sashindranath M et al (2019) Activated CD8+ T Cells Cause Long-Term Neurological Impairment after Traumatic Brain Injury in Mice. Cell Reports 29: 1178-1191.e1176 Doi 10.1016/j.celrep.2019.09.046

7 Gentleman SM, Nash MJ, Sweeting CJ, Graham DI, Roberts GW (1993) Beta-amyloid precursor protein (beta APP) as a marker for axonal injury after head injury. Neurosci Lett 160: 139–144

8 Gentleman SM, Roberts GW, Gennarelli TA, Maxwell WL, Adams JH, Kerr S, Graham DI (1995) Axonal injury: a universal consequence of fatal closed head injury? Acta neuropathologica 89: 537–543

9 Holmin S, Söderlund J, Biberfeld P, Mathiesen T (1998) Intracerebral Inflammation after Human Brain Contusion. Neurosurgery 42: 291–298 Doi 10.1097/00006123-199802000-00047

10 Johnson VE, Meaney DF, Cullen DK, Smith DH (2015) Animal models of traumatic brain injury. Handbook of clinical neurology 127: 115–128 Doi 10.1016/B978-0-444-52892-6.00008-8

11 Johnson VE, Stewart JE, Begbie FD, Trojanowski JQ, Smith DH, Stewart W (2013) Inflammation and white matter degeneration persist for years after a single traumatic brain injury. Brain 136: 28–42 Doi 10.1093/brain/aws322

12 Johnson VE, Stewart W, Arena JD, Smith DH (2017) Traumatic Brain Injury as a Trigger of Neurodegeneration. Adv Neurobiol 15: 383–400 Doi 10.1007/978-3-319-57193-5_15

13 Johnson VE, Stewart W, Smith DH (2013) Axonal pathology in traumatic brain injury. Exp Neurol 246: 35–43 Doi 10.1016/j.expneurol.2012.01.013

14 Johnson VE, Stewart W, Smith DH (2010) Traumatic brain injury and amyloid-beta pathology: a link to Alzheimer’s disease? Nat Rev Neurosci: Doi nrn2808 [pii] 10.1038/nrn2808

15 Johnson VE, Stewart W, Weber MT, Cullen DK, Siman R, Smith DH (2016) SNTF immunostaining reveals previously undetected axonal pathology in traumatic brain injury. Acta neuropathologica 131: 115–135 Doi 10.1007/s00401-015-1506-0

16 Johnson VE, Weber MT, Xiao R, Cullen DK, Meaney DF, Stewart W, Smith DH (2018) Mechanical disruption of the blood-brain barrier following experimental concussion. Acta neuropathologica 135: 711–726 Doi 10.1007/s00401-018-1824-0

17 Kilgore MD, Xiu Y, Jiang Y, Wang Y, Shi M, Zhou D, Sein T, Vodovoz SJ, Wang D, Dumont AS et al (2025) T Cell Involvement in Neuroinflammation After Traumatic Brain Injury: Implications for Therapeutic Intervention. CNS Neurosci Ther 31: e70580 Doi 10.1111/cns.70580

18 McKee CA, Lukens JR (2016) Emerging Roles for the Immune System in Traumatic Brain Injury. Front Immunol 7: 556 Doi 10.3389/fimmu.2016.00556

19 Ndode-Ekane XE, Matthiesen L, Bañuelos-Cabrera I, Palminha CAP, Pitkänen A (2018) T-cell infiltration into the perilesional cortex is long-lasting and associates with poor somatomotor recovery after experimental traumatic brain injury. Restor Neurol Neurosci Preprint: 1–17 Doi 10.3233/rnn-170811

20 Needham EJ, Helmy A, Zanier ER, Jones JL, Coles AJ, Menon DK (2019) The immunological response to traumatic brain injury. Journal of Neuroimmunology 332: 112–125 Doi 10.1016/j.jneuroim.2019.04.005

21 Needham EJ, Stoevesandt O, Thelin EP, Zetterberg H, Zanier ER, Nimer FA, Ashton NJ, Outtrim JG, Newcombe VFJ, Mousa HS et al (2021) Complex Autoantibody Responses Occur following Moderate to Severe Traumatic Brain Injury. J Immunol 207: 90–100 Doi 10.4049/jimmunol.2001309

22 Saef I, Taha Y, Omar A, Hadi A-E-H, Michael A, Tian C, Oliveira Mgd, Kuan-Jung L, Thais GM, Patrick da S et al (2025) Nasal anti-CD3 monoclonal antibody ameliorates traumatic brain injury, enhances microglial phagocytosis and reduces neuroinflammation via IL-10-dependent Treg–microglia crosstalk. Nature Neuroscience 28: 499 - 516-499-516 Doi 10.1038/s41593-025-01877-7

23 Sen T, Saha P, Gupta R, Foley LM, Jiang T, Abakumova OS, Hitchens TK, Sen N (2020) Aberrant ER Stress Induced Neuronal-IFNβ Elicits White Matter Injury Due to Microglial Activation and T-Cell Infiltration after TBI. The Journal of Neuroscience 40: 424–446 Doi 10.1523/jneurosci.0718-19.2019

24 Sherriff FE, Bridges LR, Sivaloganathan S (1994) Early detection of axonal injury after human head trauma using immunocytochemistry for beta-amyloid precursor protein. Acta Neuropathol (Berl) 87: 55–62

25 Smith DH, Nonaka M, Miller R, Leoni M, Chen XH, Alsop D, Meaney DF (2000) Immediate coma following inertial brain injury dependent on axonal damage in the brainstem. Journal of neurosurgery 93: 315–322 Doi 10.3171/jns.2000.93.2.0315

26 Tang-Schomer MD, Johnson VE, Baas PW, Stewart W, Smith DH (2012) Partial interruption of axonal transport due to microtubule breakage accounts for the formation of periodic varicosities after traumatic axonal injury. Exp Neurol 233: 364–372 Doi 10.1016/j.expneurol.2011.10.030

27 Ulyanova AV, Koch PF, Cottone C, Grovola MR, Adam CD, Browne KD, Weber MT, Russo RJ, Gagnon KG, Smith DH et al (2018) Electrophysiological Signature Reveals Laminar Structure of the Porcine Hippocampus. eNeuro 5: ENEURO.0102-0118.2018 Doi 10.1523/eneuro.0102-18.2018

28 Waltzman D, Black LI, Daugherty J, Peterson AB, Zablotsky B (2025) Prevalence of traumatic brain injury among adults and children. Ann Epidemiol 103: 40–47 Doi 10.1016/j.annepidem.2025.02.005

29 Wofford KL, Browne KD, Loane DJ, Meaney DF, Cullen DK (2024) Peripheral immune cell dysregulation following diffuse traumatic brain injury in pigs. Journal of Neuroinflammation 21: 324 Doi 10.1186/s12974-024-03317-y

30 Wolf JA, Johnson BN, Johnson VE, Putt ME, Browne KD, Mietus CJ, Brown DP, Wofford KL, Smith DH, Grady MS et al (2017) Concussion Induces Hippocampal Circuitry Disruption in Swine. J Neurotrauma: Doi 10.1089/neu.2016.4848

31 Xu L, Ye X, Wang Q, Xu B, Zhong J, Chen Yy, Wang Ll (2021) T-cell infiltration, contribution and regulation in the central nervous system post-traumatic injury. Cell Prolif 54: e13092 Doi 10.1111/cpr.13092

32 Yasir NJ, Saef I, Whalen M, McGavern D, Khoury JE (2017) Neuroimmunology of Traumatic Brain Injury: Time for a Paradigm Shift. Neuron 95: 1246-1265-1246-1265 Doi 10.1016/j.neuron.2017.07.010

